# *Pseudomonas aeruginosa* impairs mitochondrial function and metabolism during infection of corneal epithelial cells

**DOI:** 10.1101/2024.06.24.600521

**Authors:** Rajalakshmy Ayilam Ramachandran, Joelle T. Abdallah, Mahad Rehman, Hamid Baniasadi, Abigail M. Blanton, Santiago Vizcaino, Danielle M. Robertson

## Abstract

*Pseudomonas aeruginosa* (PA) is a gram-negative opportunistic pathogen that can infect the cornea as a result of trauma or contact lens wear. In addition to their known energy producing role, mitochondria are important mediators of immune signaling and host defense. While certain pathogens have developed strategies to evade host defenses by modulating host mitochondrial dynamics and metabolism, the ability of PA to harness host cell mitochondria during corneal infection is unknown. Using a combination of biochemical and imaging techniques, we show that PA infection of corneal epithelial cells induced mitochondrial fission in a DRP1-dependent manner that preceded PINK1/Parkin and FUNDC1-mediated mitophagy. PA also impaired NADH-linked respiration through a reduction in complex 1. This corresponded to a decrease in metabolic pathways related to glycolysis and the TCA cycle. Metabolomics analysis further demonstrated an upregulation of the pentose phosphate pathway, arginine, purine, and pyrimidine metabolism in PA infected cells. These pathways may provide a key source of nucleotides, amino acids, and nitrogen for both the host cell and PA, in addition to antioxidant functions. Following treatment with gentamicin to kill all extracellular bacteria, metabolic flux analysis showed that corneal epithelial cells were able to restore mitochondrial function despite the continued presence of intracellular PA. Taken together, these data demonstrate that mitochondrial dysfunction and metabolic rewiring in host cells is triggered by extracellular PA, but once inside, PA requires healthy mitochondria to ensure host cell survival.

## Introduction

*Pseudomonas aeruginosa* (PA) is a gram-negative pathogen that is associated with significant morbidity and mortality in immunocompromised patients, airway disease, and burn trauma. In the eye, PA infection can lead to blindness and when severe, evisceration or enucleation of the globe. The mechanism by which PA infects the cornea is complex and involves the formation of cholesterol enriched membrane domains known as lipid rafts that facilitate bacterial internalization.(1–4) Once inside, PA establishes an intracellular niche through the formation of membrane blebs.(5–9) Within these blebs, PA is able to survive and replicate while evading clearance by the innate immune system. Mitochondria are double membrane organelles that participate in cellular energy production via oxidative phosphorylation and the electron transport chain. Mitochondria also play important roles in regulating cell death, calcium signaling, inflammation, and cell autonomous immunity.(10–13) Mitochondria have a highly dynamic morphology that continuously changes as a result of fusion and fission events. Upon damage, mitochondria fuse to enable mixing of mitochondrial components in order to meet the metabolic demands of the cell. Mitochondrial damage can also trigger fission, a process whereby mitochondria are fragmented into smaller organelles for removal via lysosomes. The selective removal of damaged mitochondria through lysosomal degradation is known as mitophagy.(14)

Bacterial pathogens use an array of virulence factors to modify host mitochondria for their intracellular survival.(15) Bacteria such as *Listeria monocytogenes*, *Legionella pneumophilia*, and *Shigella flexneri* all induce mitochondrial fission, albeit by different mechanisms.(16–18) Rickettsia on the other hand, uses host cell mitochondrial proteins to establish an intracellular niche.(19) Many bacteria also depend upon mitochondrial function in the host cell for the generation of ATP. Such is the case for *Chlamydia trachomatis*, where the bacteria require an intact mitochondrial network to facilitate ATP uptake and intracellular bacterial survival.(20) Similar to mitochondrial fission, the effect of bacterial infection on mitophagy is varied and cell type dependent. In macrophages, *Mycobacterium sp.* have been shown to induce BCL2 interacting protein 3L (BNIP3L)-mediated mitophagy.(21) The induction of BNIP3L-mediated mitophagy in this model was associated with improved disease outcomes. Also in macrophages, *Listeria monocytogenes* is able to induce mitophagy through the NOD-like receptor, NLRX1.(16) Unlike *Mycobacterium*, *Listeria* induced mitophagy functions to buffer mtROS production by fragmented mitochondria, thereby promoting intracellular bacterial survival. *Y. Pestis* also induces mitophagy through the PTEN-induced kinase 1 (PINK1) pathway to limit mtROS.(22)

PA is capable of inducing mitophagy in worms and human embryonic kidney cells. In these cell types, mitophagy was found to be protective to host cells and prevented siderophore-mediated cell death.(23) PA has been shown to use the type 3 secretion system to target mitochondria, leading to activation of the NLRC4 inflammasome in immune cells.(24) In that model, PA simultaneously upregulated PINK1-mediated mitophagy in order to remove damaged mitochondria and limit inflammation. In lung epithelial cells, PA uses the quorum sensing molecule N-(3-oxododecanoyl)-L-homoserine lactone (3-oxo-C12-HSL) to disrupt mitochondrial architecture and impair mitochondrial metabolism.(25) Similar findings were reported in intestinal epithelial cells and fibroblasts.(26) While growing evidence suggests that PA is able to effectively manipulate host mitochondria in certain cell types, the mitochondrial response to PA infection in corneal epithelial cells is unknown.

The metabolome encompasses a diverse array of small molecules including sugars, lipids, amino acids and nucleotides that are involved in metabolic reactions. Changes in the metabolite profile in immune cells has been documented during bacterial infection.(27) In macrophages, for example, changes in host cell metabolism are associated with transition to the M1 or M2 phenotype.(28) In the cystic fibrosis airway, PA exploits the metabolites released by macrophages to increase virulence and biofilm formation.(29, 30) Despite the potentially significant role for metabolic rewiring in bacterial proliferation or containment, there is a paucity of data on the effects of PA infection on host cell metabolism in non-immune cells. In the present study, we investigated the mitochondrial and metabolic changes that occur during PA infection in corneal epithelial cells. Importantly, we show that host cells undergo distinct changes in metabolism, with an impairment in host cell mitochondria and evidence of metabolic rewiring. Upon killing extracellular PA, mitochondrial function is restored, providing a safe haven for PA to avoid clearance by the innate immune system.

## Materials and methods

### Cell culture

The human telomerase immortalized corneal epithelial (hTCEpi) cell line developed and characterized by our laboratory was used in this study.(31) Cells were maintained in defined serum-free keratinocyte growth medium (KGM) containing growth factor supplements (KGM, VWR, Randor, PA) and a 10% solution consisting of penicillin, streptomycin and amphotericin B (Lonza, Walkersville, MD). The calcium chloride concentration in KGM was maintained at 0.15 mM by supplementing additional CaCl_2_ (Calcium Chloride Solution, 0.5 M, VWR, Radnor, PA). Cells were maintained at 37^°^C and 5% CO_2_. Primary human corneal epithelial cells (HCECs) were purchased from ATCC (Mannasas, VA). The cells were maintained in defined corneal basal medium (ATCC) and supplemented with the corneal epithelial cell growth kit (ATCC) and Penicillin-Streptomycin-Amphotericin B.

### Bacterial strain

A standard invasive strain of *Pseudomonas aeruginosa*, PAO1, was used in these studies (generous gift from Dr. Suzanne M J Fleiszig, University of California, Berkeley). Bacteria were maintained on tryptic soy agar (TSA, Fisher Scientific, Waltham, MA). For infection assays, PA was grown to log growth phase on a tryptic soy agar slant. PA was resuspended in PBS and adjusted to an OD of ∼1 X 10^8^ CFU/mL. The concentration of PA was confirmed by serial dilution in PBS and plating on tryptic soy agar for colony count determination. Plates were incubated overnight at 37°C. For all experiments, hTCEpi cells were infected at a concentration of 10 bacteria per cell. With the exception of the time course studies to monitor the expression and phosphorylation of mitochondrial fission proteins, a 2 hour infection time point was used for all experiments. For the metabolic flux studies, heat killed PA was used as an additional control. PA were heat killed by incubation at 80°C for 30 minutes. Heat killed bacteria were then serially diluted in PBS and plated on tryptic soy agar to confirm that all PA were non-viable.

### Western blot

Cell lysates were collected in RIPA buffer (Thermo Fisher, Rockford, IL) containing a protease and phosphatase inhibitor cocktail (Thermo Fisher, Rockford, IL). Samples were mixed with 4X lamelli buffer (Bio-rad, Hercules, CA) and separated based on molecular weight by running SDS-PAGE. The separated proteins were blotted onto PVDF membranes (Bio-rad, Hercules, CA), blocked with 5% milk in TBST (Bio-rad, Hercules, CA), and then incubated with one of the following primary antibodies overnight at 4°C: PINK1 (#6946), FUNDC1 (#49241), Parkin (#3283), BNIP3/NIX (# 12396), FIS1 (#32525), DRP1 (#8570), pDRP1 Ser616 (#3458), MFN1 (#14739), MFN2 (#11925), OPA1 (#67589), Vinculin-HRP (#18799) and β-actin-HRP (#47778) (Cell Signaling, Danvers, MA); and OXPHOS (#45-8199, Thermo Fisher, Rockford, IL). For primary antibodies without an HRP tag, membranes were incubated in an appropriate secondary antibody: anti-Rabbit IgG-HRP Conjugate #1706515 and anti-Mouse IgG (H + L)-HRP Conjugate #1706516 (Bio-Rad, Hercules, CA). Image Quant TL Toolbox v8.1 software (Amersham Bioscience, Piscataway, NJ) was used for image quantification. Protein bands were normalized to Vinculin or β-actin (internal control). Data is shown as fold change in protein in PA infected cells compared to the control. For phosphorylated proteins, data is shown as the ratio of internal control normalized phosphorylated protein to internal control normalized non-phosphorylated protein.

### JC-1 staining

hTCEpi cells were infected with PAO1 for 2 hours. Ten minutes prior to the end of the incubation period, cells were treated with 10 μg/ml of tetraethylbenzimidazolyl-carbocyanine iodide (JC-1) dye (Invitrogen/Molecular Probes, Eugene, OR). As a positive control, hTCEpi cells were treated with 50 μM carbonyl cyanide m-chlorophenyl hydrazine (FCCP) for 10 minutes at 37^°^C. Cells were washed 3 times with PBS and imaged using a Leica SP8 laser scanning confocal microscope (Leica Microsystems, Heidelberg, Germany) using a 63x water objective. The microscope was enclosed within an environmental chamber (Life Imaging Services, Basel, Switzerland) to maintain cells at 5% CO2, 37°C during imaging. JC-1 monomers were captured using the 488 nm excitation laser and JC-1 aggregates were captured using the 561 nm excitation laser. Cells were sequentially scanned to avoid any spectral crosstalk. Image J was used to measure the ratio of red to green fluorescence in the respective images.(32)

### Transmission electron microscopy

hTCEpi cells were seeded onto 35 mm coverslip bottom dishes (MatTek, Ashland, MA) and infected with PAO1 for 2 hours. At the end of the incubation period, cells were fixed for 15 minutes at room temperature using 2.5% glutaraldehyde/0.1 M cacodylate buffer (pH 7.4). Cells were then rinsed several times with 0.1 M sodium cacodylate buffer, incubated with 1% osmium tetroxide + 0.8% K3[Fe(CN6)] in 0.1 M sodium cacodylate buffer for 1 hour at room temperature and stained overnight with 2% aqueous uranyl acetate. Stained cells were dehydrated with ethanol, infiltrated with Embed-812 resin, and polymerized at 60°C overnight. Sections were cut using a diamond knife (Diatome US, Quakertown, PA) on a Leica Ultracut UCT (7) ultramicrotome (Leica Microsystems, Wetzlar, Germany). After placing sections onto copper grids, the samples were post-stained with aqueous 2% uranyl acetate and lead citrate. The images were captured using a JEOL 1400 Plus (JEOL, Peabody, MA) equipped with a LaB6 source with 120 kV.

### Metabolic flux assay

A Seahorse Metabolic Flux Analyzer XFp (Agilent Technologies, Santa Clara, CA) was used to measure the cellular oxygen consumption rate (OCR), non-mitochondrial respiration, maximal respiration, and spare respiratory capacity. An XFp Cell Mito Stress Test Kit (Agilent Technologies, Santa Clara, CA) was used per the manufacturer’s instructions. In brief, cells were seeded onto Seahorse XFp mini plates and infected with bacteria for 2 hours. At the end of the incubation period, the cells were treated with 200 μg/mL of gentamicin (Sigma, St. Louis, MO) in Seahorse XFp medium (Agilent Technologies, Santa Clara, CA) for 30 minutes to kill all extracellular PA. For methyl-β-cyclodextrin (MβCD) (MedChemExpress, Monmouth Junction, NJ) treatment, cells were pre-treated with 5 mM MβCD in KGM for 1 hour, followed by a 2 hour incubation of PAO1 in KGM containing 5 mM MβCD. Finally, cells were incubated for 30 minutes in 5 mM MβCD in fresh XFp medium (Agilent Technologies, Santa Clara, CA). To interrogate mitochondrial function, 10 μM oligomycin was injected in to the system at 20 minutes to inhibit ATP synthase. At 50 minutes, 10 μM FCCP was injected to uncouple oxidative phosphorylation from the electron transport chain, allowing for maximum electron flow. Finally, rotenone and actinomycin A were used to inhibit mitochondrial complex I and III, respectively. Data were recorded every 6 minutes, for a total of 94 minutes. Wave software, version 2.3.0 (Agilent Technologies, Santa Clara, CA) was used for data analysis. Data was normalized using total cell number per plate. Cell numbers were quantified using a Celigo Imaging Cytometer (Nexelom Bioscience, Lawrence, MA). The following equation was used for calculating spare respiratory capacity: spare respiratory capacity = (maximal respiration)/(basal respiration) × 100.

### Gentamicin Protection Assay

A gentamicin protection assay was used to measure PAO1 internalization in hTCEpi cells. Cells were seeded at 50-60% confluence and allowed to adhere overnight. Culture media was changed the following morning. Cells were then infected with PAO1 at a cell:bacteria ratio of 1:500. After 2 hours, cells were washed 3 times in PBS and treated with 200 µg/mL gentamicin in culture media at 37°C for 1 hour to kill all extracellular bacteria. Cells were then washed three times in PBS and lysed with 0.25% TritonX-100 in PBS for 15 minutes. All samples were serially diluted and plated in triplicate on trypic soy agar for colony count determination.

### Metabolomics

hTCEpi cells were infected with overnight culture of PAO1 and metabolites were extracted in 80% methanol with 5 μM heavy internal standard.(33) In brief, at the end of the treatment period, cell culture supernatants were removed and ice cold 80% methanol with 5 μM heavy internal standard was added onto the top of the cells. Cells were then kept on ice for 5 minutes, scrapped, and collected along with 80% methanol. The samples were freeze thawed 1 minute each in dry ice and 4^°^C for 5 times to collect metabolites, and then centrifuged to remove the pellet. The supernatant was filtered using 0.2 µM centrifugal filters and LC-MS was performed in Sciex QTRAP 6500+ mass spectrometer with EI ion spray ion in both positive and negative mode. A Shimadzu HPLC (Nexera X2 LC-30AD) with analyst 1.7.2 software was coupled to the mass spectrometer. A SeQuant® ZIC®-pHILIC 5 μm polymeric 150 × 2.1 mm PEEK coated HPLC column with a temperature of 45^°^C, injection volume of 5 µL and flow rate of 0.15 mL/min was used. The mobile phase was solvent A: acetonitrile and solvent B: 20 mM ammonium carbonate with 0.1% ammonium hydroxide and 5 µM of medronic acid. The gradient elution procedure was as follows: 0 min: 80% B, 20 min: 20% B, 20.5 min 80% B, 34 min: 80% for a total of 34 minutes. SCIEX MultiQuant 3.0.3 software was used for the detection and quantification of metabolites and MetaboAnalyst 5.0 software was used for data representation. SCIEX MultiQuant 3.0.3 software was used for the detection and relative quantification of metabolites based on peak area. The peak area of each feature was normalized to the internal standard. MetaboAnalyst 5.0 (McGill University, Montreal, Canada) was used for statistical analysis and data representation. An adjusted p-value < 0.05 was considered significant. For pathway analysis, a false discovery rate of <0.1 was considered significant.

## Results

### PA infection induces mitophagy and mitochondrial membrane depolarization in corneal epithelial cells

Mitophagy is a major mitochondrial quality control mechanism in mammalian cells. To investigate the effects of PA infection on mitophagy, hTCEpi cell lysates were subject to western blotting for mitophagy proteins PINK1, Parkin, FUNDC1, and BNIP3L/NIX (Figure 1 A-H). Two hours post infection, there was a significant increase in the expression of mitophagy proteins PINK1, Parkin, and FUNDC1 compared to non-infected control cells. In contrast to this, the expression of BNIP3L/NIX was reduced. Since PINK1 is involved in mitophagy associated with mitochondrial membrane depolarization, we next quantified mitochondrial membrane potential using JC-1, a cationic probe that accumulates in mitochondria (Figure 1, I-J). In non-infected control cells, we observed red fluorescence interspersed throughout green fluorescing mitochondria, indicating the formation of JC-1 aggregates in highly polarized mitochondria. In addition, mitochondria were arranged in a tubular network. In PA infected cells, both red and green fluorescence were reduced. Mitochondria lost their elongated tubular structure, shifting towards a more condensed peri-nuclear pattern. Quantification of the ratio of red to green fluorescence confirmed a significant reduction in mitochondrial polarization during infection (Figure 1J). Similar findings were observed in primary cultured HCECs (Supporting figure 1A). Together, these findings demonstrate that PA infection triggers mitochondrial membrane depolarization, followed by the induction of PINK1/Parkin and FUNDC1-mediated mitophagy in corneal epithelial cells.

**Figure 1:**
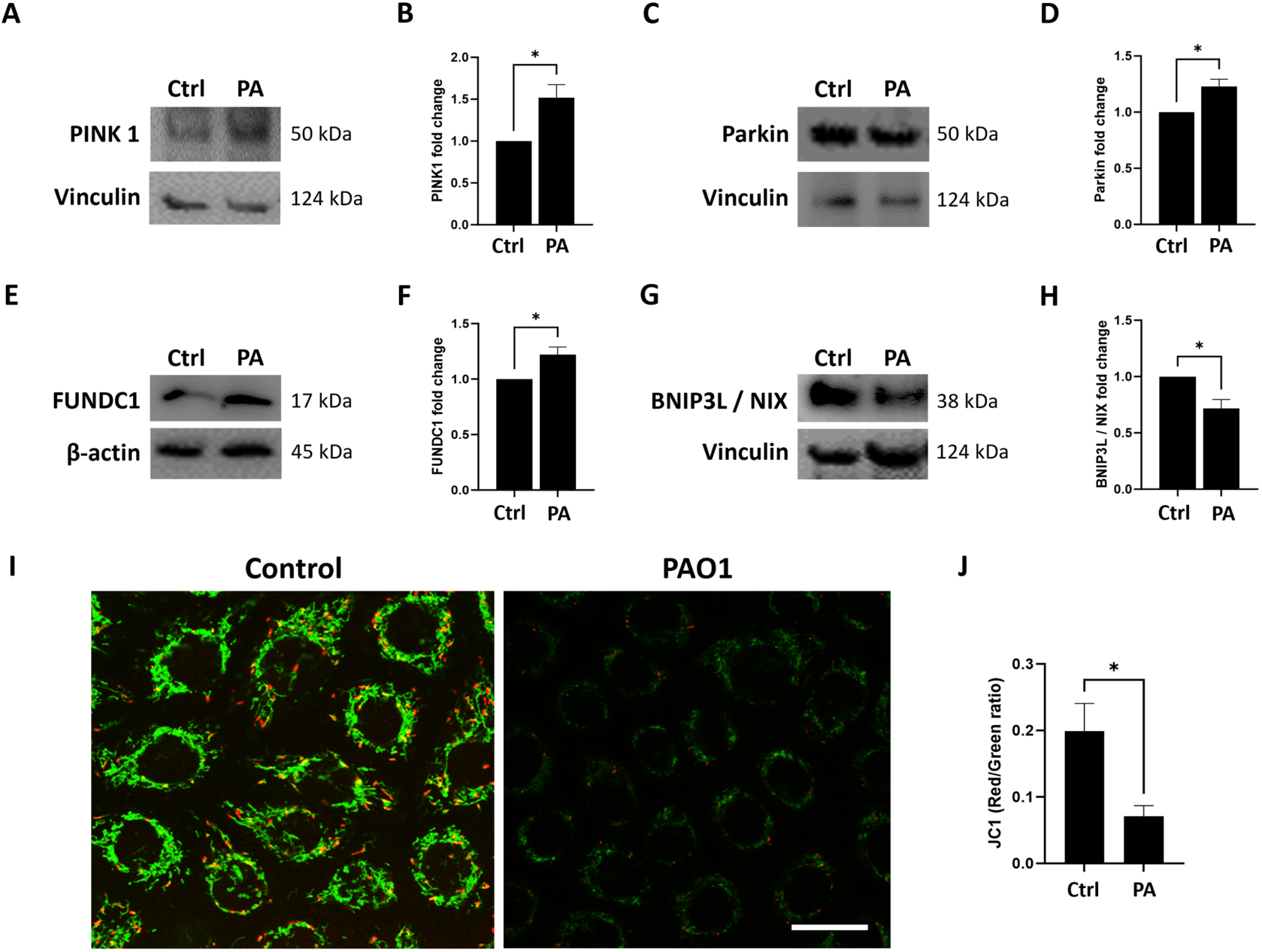
PA infection induces mitophagy and mitochondrial depolarization in corneal epithelial cells. Western blot was performed to detect mitophagy proteins and JC-1 staining was used as an indicator of mitochondrial membrane potential. (A-H) Representative western blot and quantitative densitometry of respective mitophagy proteins normalized to vinculin or β-actin. PA infection of hTCEpi cells increased (A-B) PINK1, (C-D) Parkin, and (E-F) FUNDC1 expression compared to the uninfected control cells. (G-H) BNIP3L/NIX expression was downregulated. (I) JC-1 monomers shown in green, and aggregates in red. (J) Quantification of the red to green ratio indicates that PAO1 infection induced mitochondrial depolarization. Data normalized to the non-infected control, N≥3, *P<0.05, students t-test. Images representative of 3 repeated experiments.

### PA infection induces mitochondrial fission in corneal epithelial cells

Upon damage, mitochondria first undergo fission in preparation for mitophagy. To confirm that PA infection induced fission in corneal epithelial cells, the expression level of FIS1 and phosphorylation of DRP1 at ser616 were examined over time. As early as 5 minutes post infection, we found a significant increase in pDRP1 (Figure 2, A-B). This increase was maintained at 15 minutes, leveling off 30 minutes post infection (Figure 2, C-F). At the 2 hour time point, the level of pDRP1 was reduced compared to the non-infected control (Figure 2 G-H). Like pDRP1, FIS1 was also increased at 5 and 15 minutes post infection, again leveling off at 30 minutes (Figure 2, I-N). At 2 hours however, there was a secondary increase in FIS1 expression (Figure 2, O-P). Collectively, these data confirm the induction of mitochondrial fission following PA infection.

**Figure 2:**
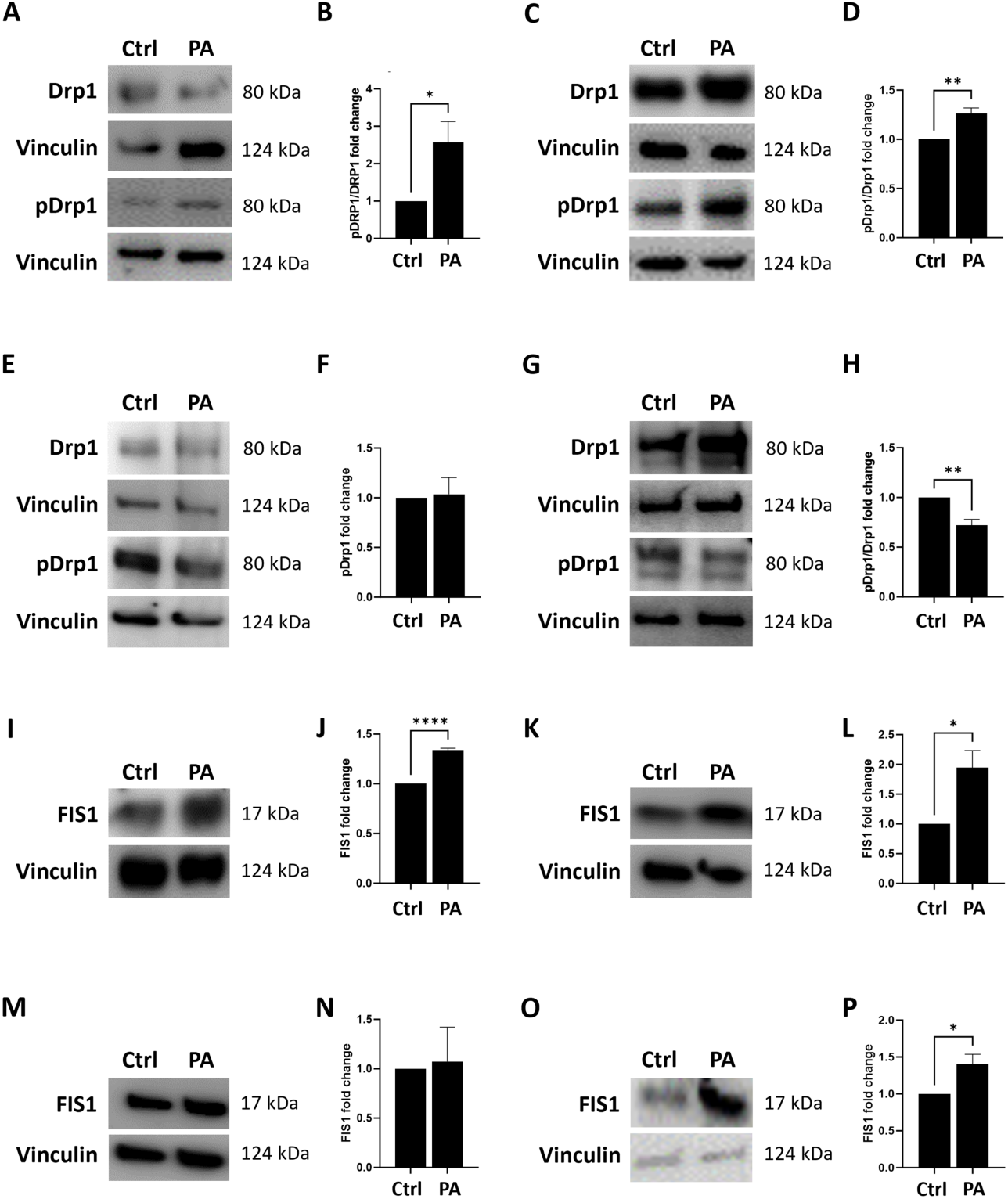
PA infection induces mitochondrial fission in corneal epithelial cells. (A-H) Western blot was performed to examine mitochondrial fission proteins at the indicated time points post infection. Phosphorylated DRP1 (ser616 residue) was normalized to total DRP1. Representative western blot and densitometry measurements at (A-B) 5 minutes, (C-D) 15 minutes, (E-F) 30 minutes, and (G-H) 2 hours post infection in hTCEpi cells. (I-P) Representative western blot and densitometry measurements of FIS1 normalized to vinculin at (I-J) 5 minutes, (K-L) 15 minutes, (M-N) 30 minutes, and (O-P) 2 hours post PA infection in hTCEpi cells. Blots show an increase in mitochondrial fission at 5 and 15 minutes after PA infection. Data normalized to non-infected control cells, N≥3, *<0.05, **p<0.01, ****p<0.0001, students t-test.

### PA infection rewires corneal epithelial cell metabolism during infection

To gain further insight into the metabolic changes in corneal epithelial cells during infection, we performed metabolomics. Metabolites were extracted from PA infected and non-infected corneal epithelial cells and subject to LC-MS using an untargeted approach. A total of 164 metabolites were detected. Principal component analysis (PCA) and clustering was performed for all 164 metabolites to see the overall differences between groups. The PCA score plot, shown in Figure 3A, demonstrates distinct metabolite profiles between infected and non-infected cells. Similarly, clustering results also show that the two groups were clustered into separate nodes (Figure 3B), indicating a shift in the corneal epithelial cell metabolome during PA infection. A total of 34 significantly different host metabolites (fold change ≥ 2 or ≤ -2) were detected and are plotted as a heat map (Figure 3C). Of these, 25 metabolites were upregulated and 8 were downregulated. Pathway enrichment analysis of the 34 metabolites is shown in Supporting figure 2 (A-C). The most enriched metabolic pathways included *de novo* triacylglycerol biosynthesis, betaine metabolism, cardiolipin biosynthesis, the glycerol phosphate shuttle, and purine metabolism. Biotin metabolism was the most downregulated pathway. We next performed pathway impact analysis (Figure 4, A-C). The most significant and highly impactful pathways that were increased in host cells during infection included arginine biosynthesis, purine and pyrimidine metabolism, and the pentose phosphate pathway. Downregulated pathways included pyruvate metabolism, alanine, aspartate and glutamate metabolism, nicotine and nicotinamide metabolism, glycolysis/gluconeogenesis, and the TCA cycle. Overall, these data indicate a reduction in the metabolic pathways related to energy generation in host cells.

**Figure 3:**
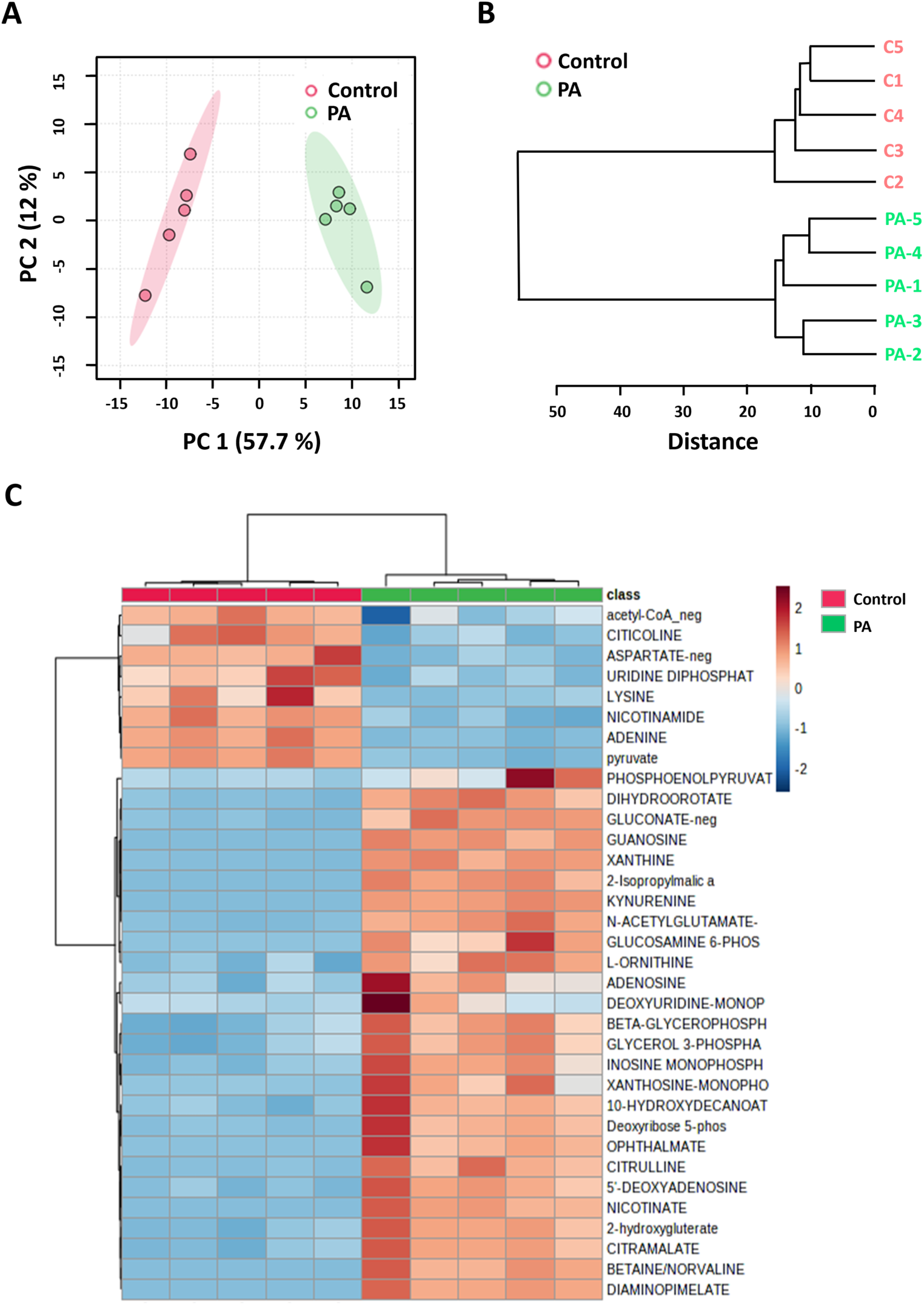
PA infection induces a unique metabolite signature in the corneal epithelium. (A) PCA score plots of the total metabolites detected in samples. (B) Hierarchical clustering dendrogram of control and PA groups. (C) Heat map of 34 significant metabolites between the groups. Untargeted metabolomics N=5, MetaboAnalyst 5.0.

**Figure 4:**
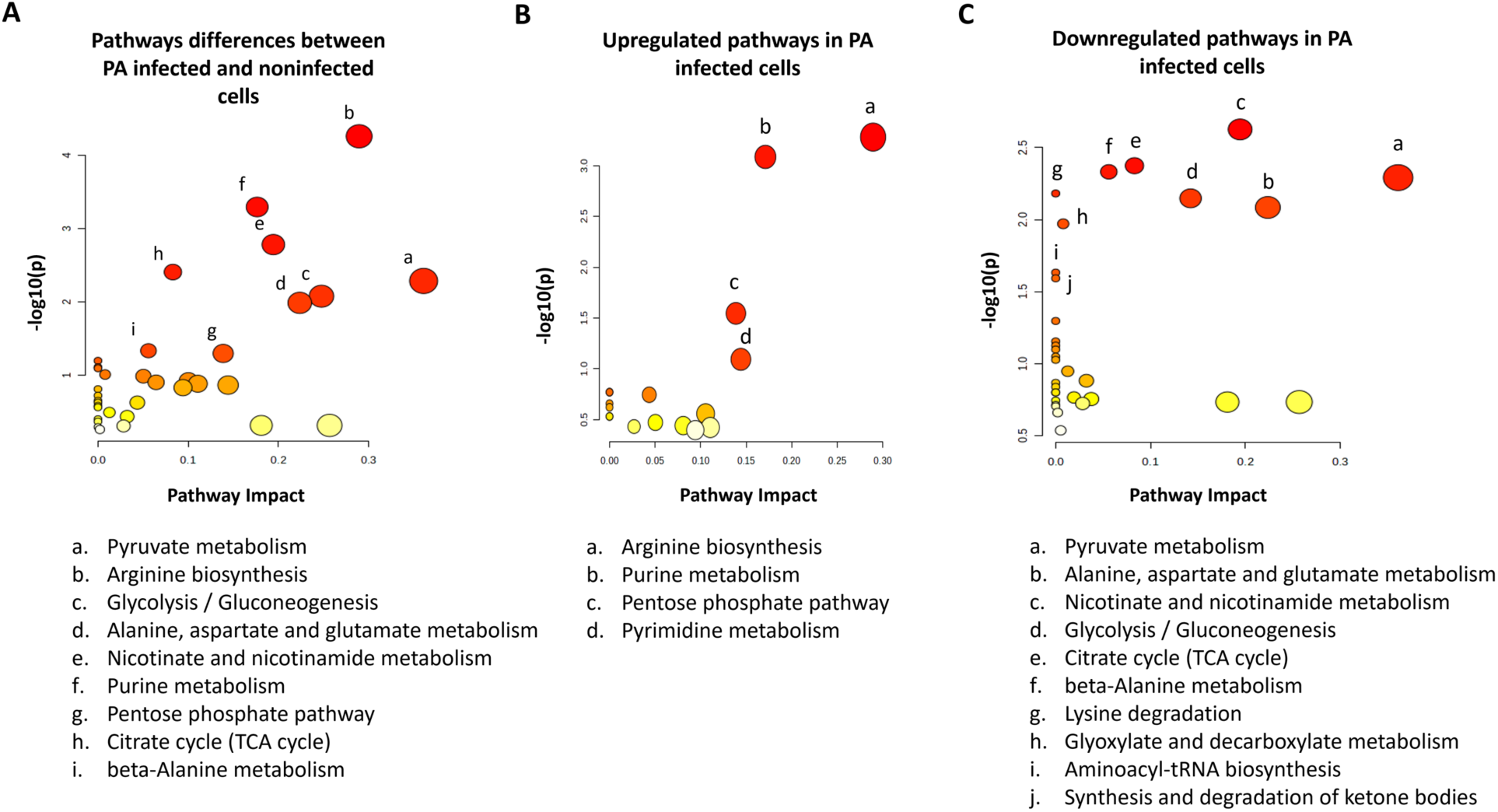
PA rewires corneal epithelial cell metabolism by downregulating the TCA cycle and glycolysis; and increasing the pentose phosphate pathway. (A) Pathway impact plot showing the most significant pathways related to 34 significantly different metabolites between non-infected and PA infected groups. (B) Pathway impact plot showing the most significant pathways related to 26 significantly upregulated metabolites in the PA infected group. (C) Pathway impact plot showing the most significant pathways related to 8 significantly downregulated metabolites in the PA infected group. Untargeted metabolomics N=5, MetaboAnalyst 5.0.

### PA infection alters mitochondrial respiration in corneal epithelial cells

Due to the changes observed using metabolomics, we next evaluated the effects of PA infection on mitochondrial respiration by western blotting for proteins involved in oxidative phosphorylation (Figure 5, A-E). Compared to control cells, there was a measurable decrease in complex I expression in PA infected cells. This indicates a decrease in NADH-linked respiration. In contrast to this, PA infection had no effect on complexes III, IV or V. We next measured mitochondrial respiration using a Seahorse metabolic flux assay. The oxygen consumption rate (OCR) was plotted as a function of time for PA infected and non-infected hTCEpi cells (Figure 6A). For these experiments, heat killed (HK) PA was used as an additional control. In terms of the total OCR (mitochondrial OCR + non-mitochondrial OCR), there were no differences between cells treated with HK PA and control cells (Figure 6, A-B). In contrast to this, the addition of viable PA to cells increased total OCR twofold. This corresponded to a large increase in non-mitochondrial OCR (Figure 6C) in PA infected cells. This large increase in non-mitochondrial OCR represents a combination of oxygen consumption by viable PA and non-mitochondrial oxygen consumption by host cells. HK PA also increased non-mitochondrial respiration; however, this effect was small in magnitude, with the signal coming entirely from host cells. Following treatment with oligomycin to inhibit ATP synthase and FCCP to uncouple the electron transport chain from oxidative phosphorylation, thereby allowing for maximum electron flow across the membrane, maximum and spare respiratory capacity were calculated. There was a complete loss of maximum and spare respiratory capacity in hTCEpi cells infected PA (Figure 6, D-E). In cells treated with HK PA, both maximal and spare respiratory capacity were decreased compared to control cells, but greater than that seen for PA infected cells. Since spare respiratory capacity is an important factor in cell health, the drop in response to HK PA indicates that even non-viable PA are able to induce a measurable level of stress in host cells.

**Figure 5:**
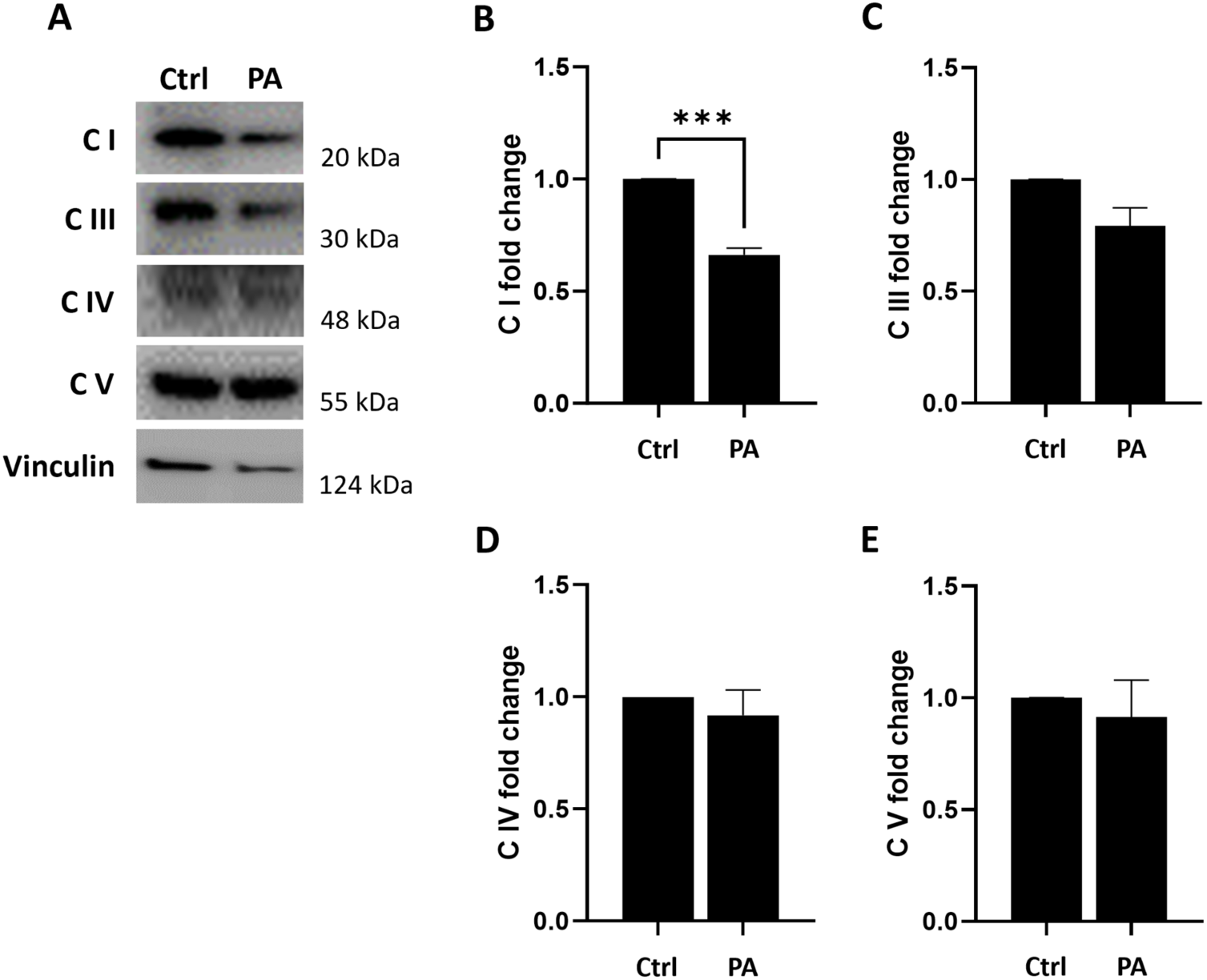
PA targets OXPHOS complex I to alter mitochondrial respiration. Western blot was performed to detect OXPHOS complex I, III, IV, and V proteins 2 hours post infection. (A-E). Representative western blot and quantitative densitometry measurement of respective proteins are shown for (B) complex I, (C) complex III, (D) complex IV, and (E) complex V. Data normalized to the non-infected control. N≥3, ***P< 0.001, students t-test.

**Figure 6:**
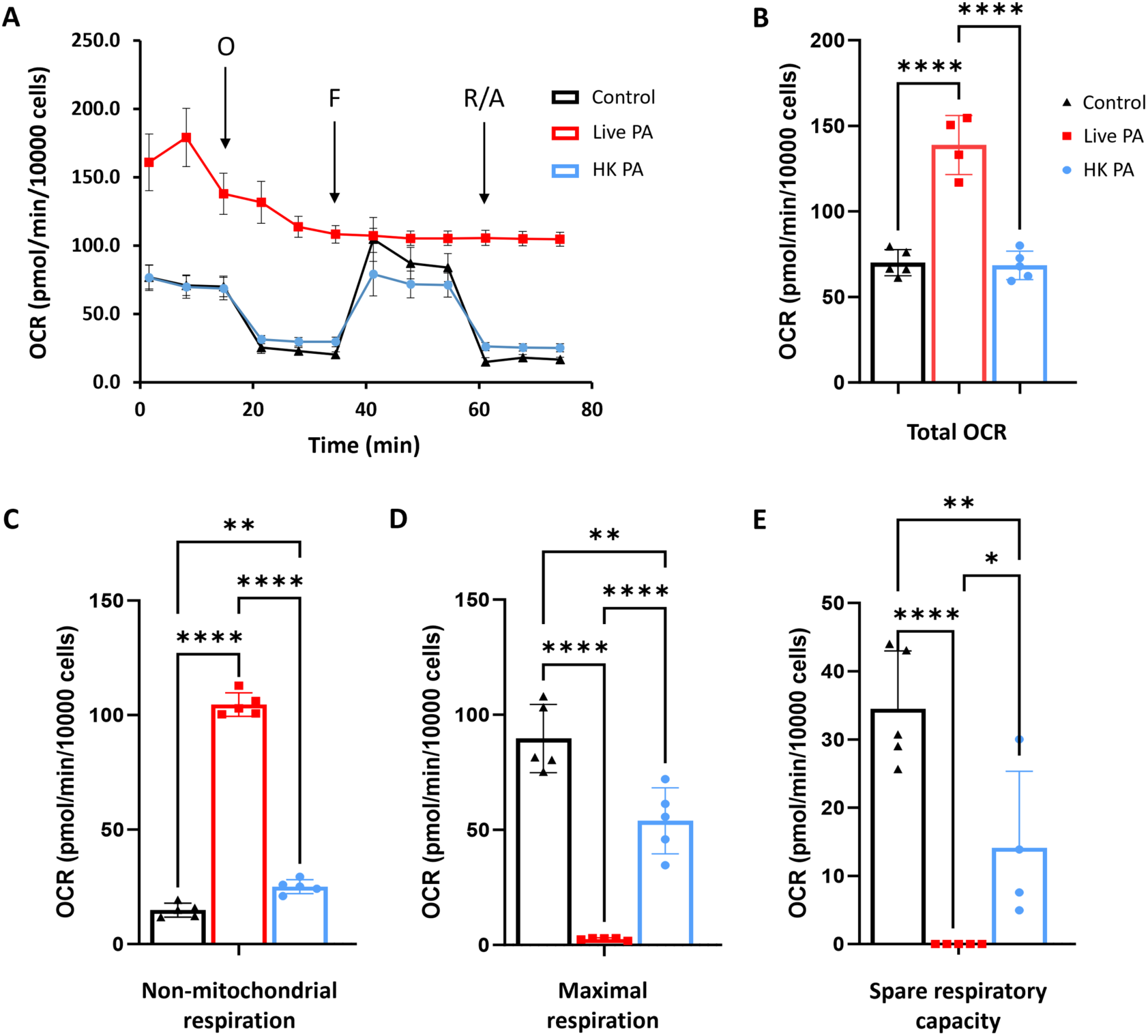
Viable and heat-killed PA reduce spare respiratory capacity in corneal epithelial cells. Two hours post infection, mitochondrial respiration was assessed using a Seahorse metabolic flux assay. (A) The oxygen consumption rate (OCR) was plotted as a function of time for all groups. (B) Total OCR (basal OCR + non-mitochondrial O_2_ consumption). (C) Non-mitochondrial oxygen consumption. (E) Maximal respiration. (E) Spare respiratory capacity. There was an increase in overall oxygen consumption in cells treated with PA that corresponded to an increase in non-mitochondrial oxygen consumption. HK = heat killed PAO1. Data normalized to total cell number, N≥3, *P<0.05, **P<0.01, ****P<0.0001. One-way ANOVA, Tukey’s post hoc multiple comparison test.

### Extracellular, not intracellular, PA alters mitochondrial respiration during infection

To determine the effect of extracellular versus intracellular PA on mitochondrial respiration, hTCEpi cells infected with PA were treated with gentamicin to kill all extracellular bacteria prior to the Seahorse assay. The presence of intracellular PA was confirmed using a gentamicin protection assay (Supplementary figure 3). As expected, the presence of viable extracellular PA showed a large increase in total OCR and non-mitochondrial OCR (Figure 7, A-C). However, in gentamicin treated cells that retained intracellular PA, total OCR and non-mitochondrial OCR were similar to non-infected controls. This change in OCR suggests that PA may be shifting from aerobic to anaerobic metabolism when it moves from the extracellular to the intracellular environment. Surprisingly, the presence of intracellular PA had no adverse effect on maximum or spare respiratory capacity (Figure 7, D-E). Taken together, these data suggest that once PA becomes intracellular, host cell mitochondria are able to rapidly restore function.

**Figure 7:**
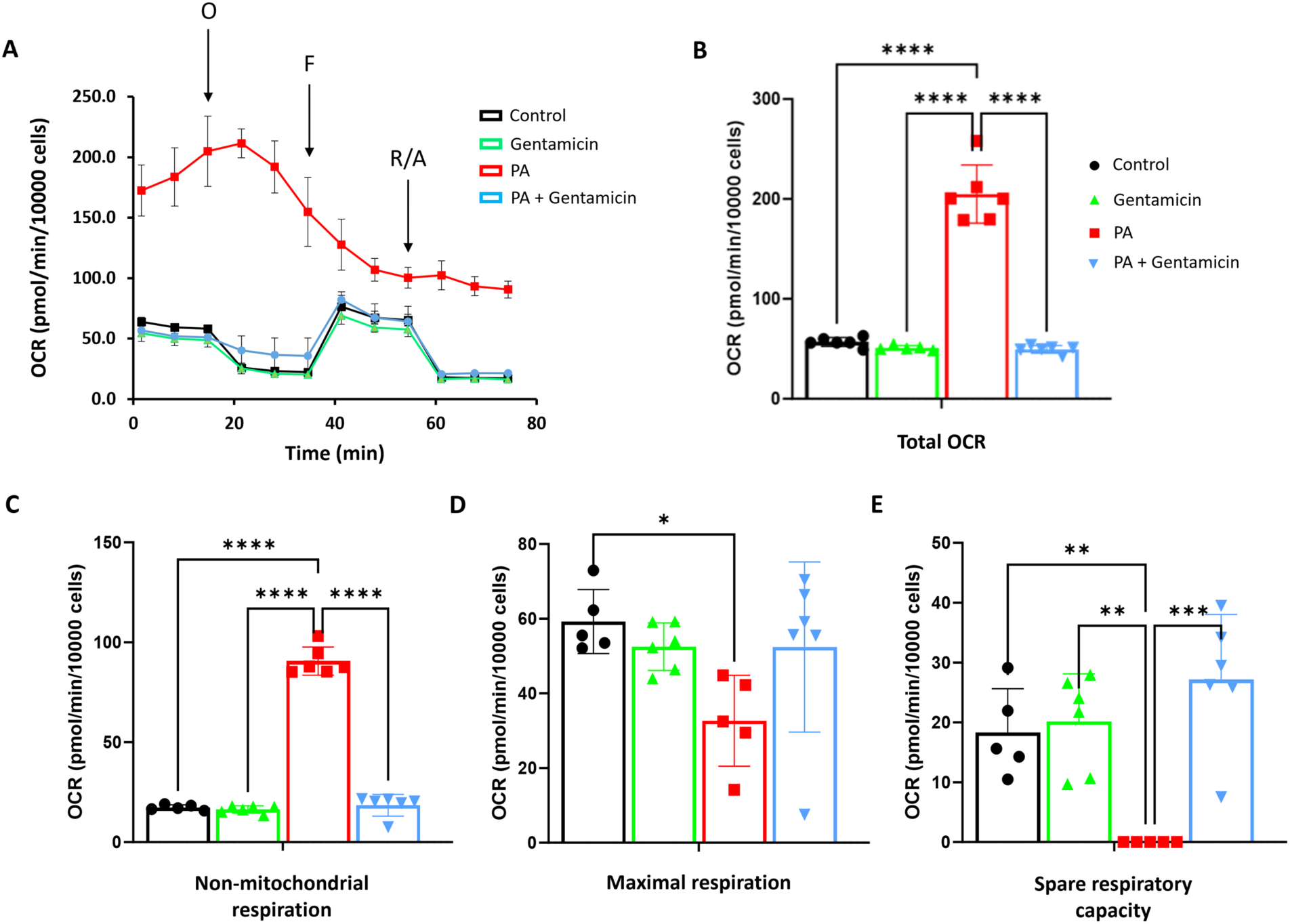
Host cell mitochondrial function is restored following gentamicin treatment. PA-infected cells were treated with gentamicin to kill all extracellular bacteria. Mitochondrial respiration was then assessed using a Seahorse metabolic flux assay. (A) The oxygen consumption rate (OCR) was plotted as a function of time for all groups. (B) Total OCR (basal OCR + non-mitochondrial O_2_ consumption). (C) Non-mitochondrial oxygen consumption. (E) Maximal respiration. (E) Spare respiratory capacity. Treatment with gentamicin restored metabolic parameters to control levels. Data are normalized to total cell number, N≥3, *P<0.05, **P<0.01, ***P<0.001, ****P<0.001. One-way ANOVA, Tukey’s post hoc multiple comparison test.

### Disruption of lipid-raft mediated PA internalization increases spare respiratory capacity

Previous work has shown that PA undergoes lipid raft-mediated internalization in corneal epithelial cells. Methyl-beta-cyclodextrin (MβCD), a lipid raft inhibitor that disrupts raft function by depleting cholesterol from cell membranes, has been shown to reduce the ability of PA to invade hTCEpi cells.(1) To further investigate the role of intracellular PA on hTCEpi cell metabolism, we first pre-treated cells with MβCD, as reported by our laboratory group.(1) In PA infected cells, MβCD decreased the total OCR compared to PA infected cells (Figure 8, A-B). However, it remained increased compared to the non-treated cells. In contrast, MβCD was not able to blunt the PA-mediated increase in non-mitochondrial OCR (Figure 8C). Unexpectedly, MβCD had a significant effect on maximal respiration and spare respiratory capacity, despite the presence of viable extracellular PA (Figure 8, D-E). In PA infected cells, MβCD not only recovered both parameters, but increased them twofold over control levels. The cause of this increase in spare respiratory capacity in unknown. While MβCD can be used to deplete cholesterol from both the plasma and mitochondrial membranes, which could affect mitochondrial function, treatment of non-infected hTCEpi cells with MβCD failed to increase spare respiratory capacity.

**Figure 8:**
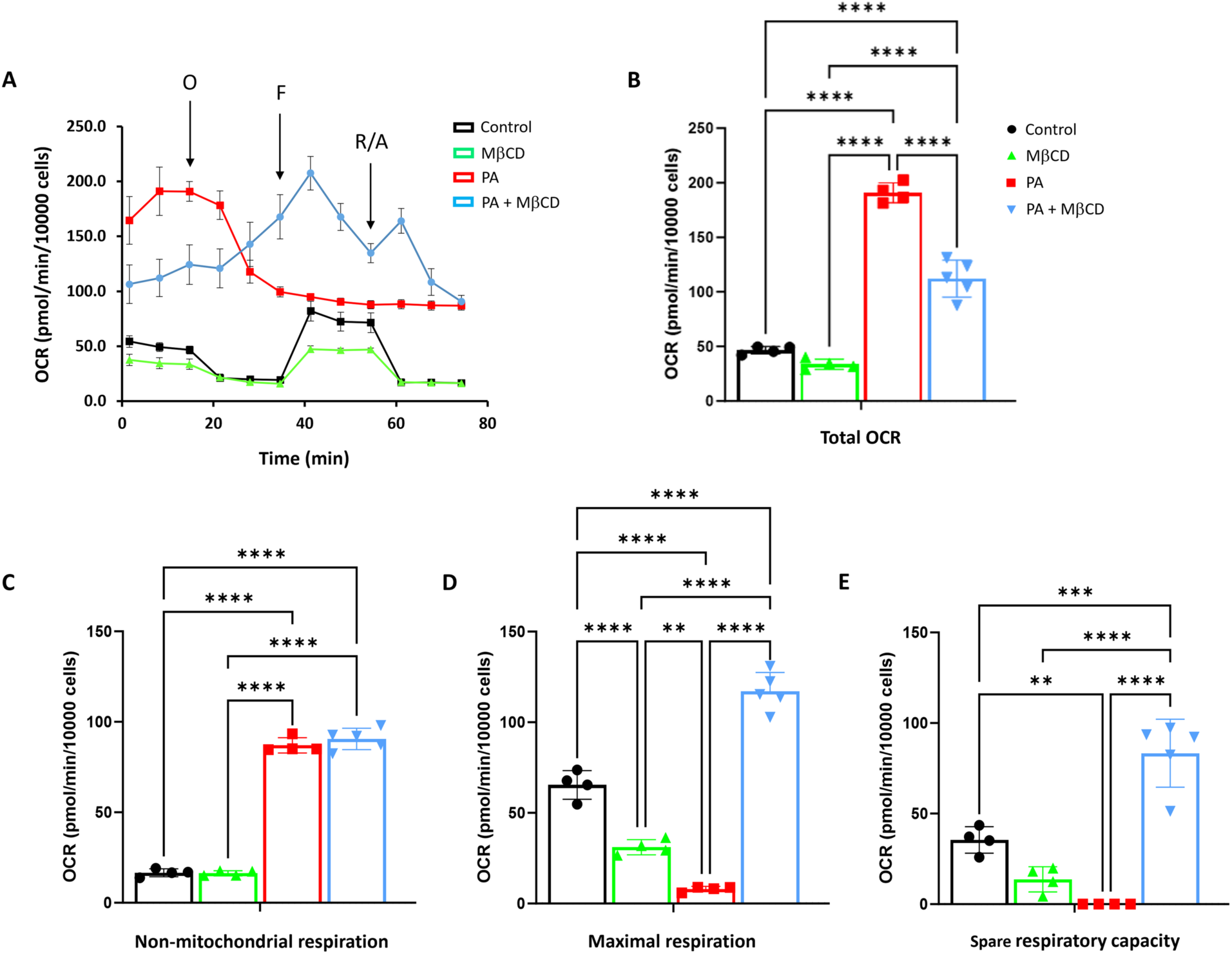
Disruption of lipid raft-mediated PA internalization increases spare respiratory capacity. hTCEpi cells were treated with the lipid raft inhibitor methyl-beta-cyclodextrin (MβCD) prior to PA infection. Cells were then treated with PA for 2 hours. Mitochondrial respiration was assessed using a Seahorse metabolic flux assay. (A) The oxygen consumption rate (OCR) was plotted as a function of time for all groups. (B) Total OCR (basal OCR + non-mitochondrial O_2_ consumption). (C) Non-mitochondrial oxygen consumption. (E) Maximal respiration. (E) Spare respiratory capacity. Treatment with MβCD increased maximal and spare respiratory capacity. Data are normalized to total cell number, N≥3, **P<0.01, ***P<0.001, ****P<0.001. One-way ANOVA, Tukey’s post hoc multiple comparison test.

### Electron microscopy confirms mitochondrial fragmentation during PA infection

To further investigate the effects of gentamicin and MβCD on mitochondria, we next evaluated mitochondrial ultrastructure using transmission electron microscopy (TEM, Figure 9). Mitochondria in control, non-infected hTCEpi cells were elongated with normal, lamellar cristae (Figure 9A). Consistent with our western blotting data suggesting an increase in fission, mitochondria in PA infected cells were small and round with wavy or missing cristae. In primary cultured HCECs, control cells displayed normal, elongated mitochondria similar to hTCEpi cells (Figure 9B). In PA infected HCECs, we again observed many small, round, mitochondria with few cristae. In non-infected cells, mitochondria were of average length and retained intact lamellar cristae (Figure 9C). In PA infected cells treated with gentamicin, there was no evidence of mitochondrial fission. Instead, gentamicin treatment prevented mitochondrial fission in PA infected cells. Since MβCD can affect cholesterol depletion across all cell membranes, we also examined the effects of MβCD on mitochondrial ultrastructure (Figure 9D). While gentamicin treatment was able to block mitochondrial fission due to the removal of extracellular PA, MβCD had no observable effect on mitochondrial dynamics. Instead, treatment with MβCD enhanced lamellar cristae architecture. This increase in cristae architecture may explain, in part, the observed increase in spare respiratory capacity. Taken together, these data show that mitochondria in corneal epithelial cells are able to adapt and regenerate despite the presence of intracellular PA without a corresponding effect on mitochondrial dynamics. A schematic of our overall findings is shown in Figure 10.

**Figure 9:**
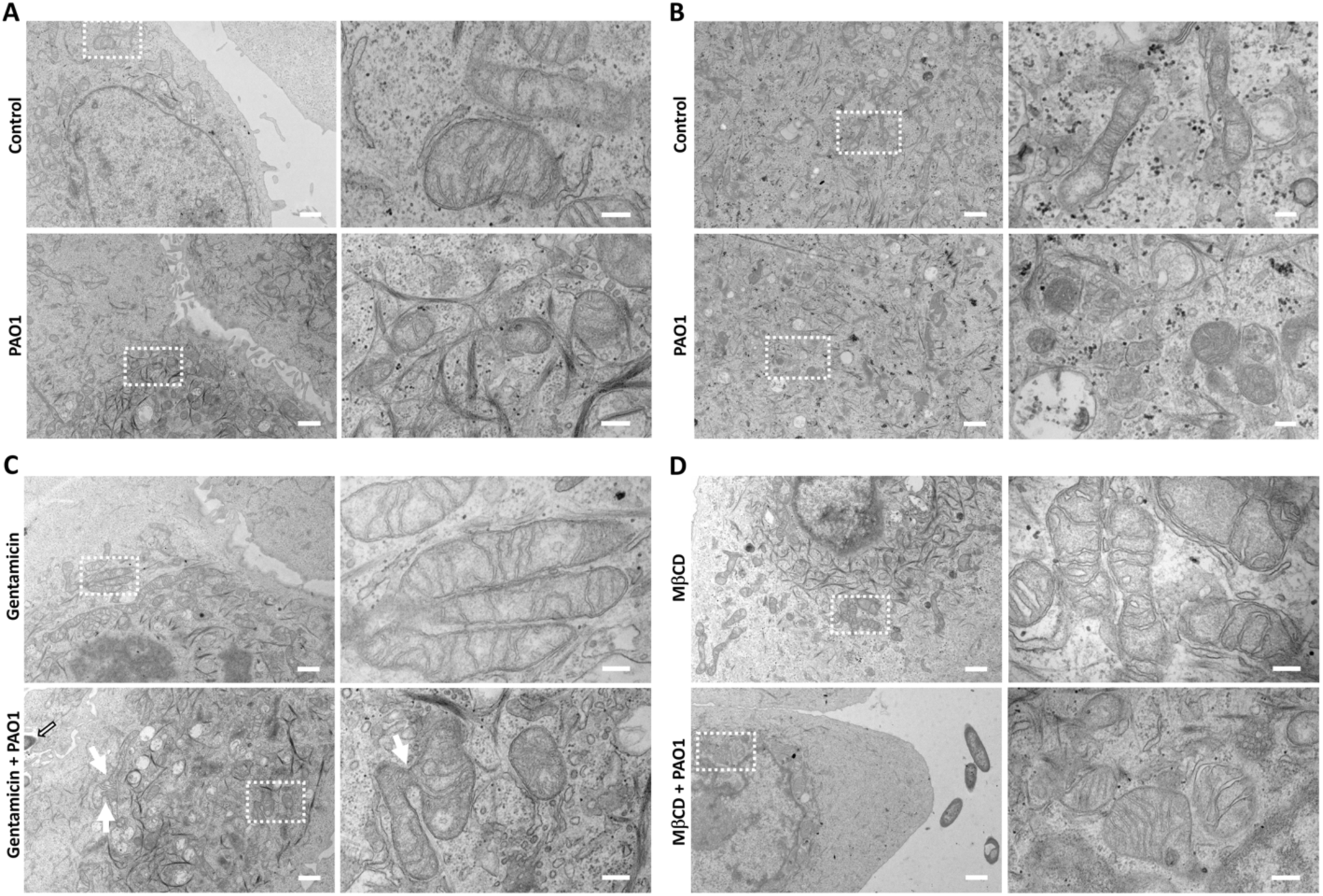
Gentamicin and MβCD both induce changes in mitochondrial ultrastructure. (A) TEM of control and PA-infected hTCEpi cells at 2 hours post infection. (B) TEM of control and PA-infected HCECs at 2 hours post infection. In both cell types, mitochondria in PA infected cells were small and round, with a loss of lamellar cristae architecture. (C) TEM of hTCEpi cells treated with gentamicin to kill extracellular PA. Gentamicin treatment of PA infected cells showed evidence of mitochondrial hyperfusion (white arrows) with partial restoration of lamellar cristae morphology. Open arrows indicate intracellular PA. (D) TEM of hTCEpi cells treated with the lipid raft inhibitor, methyl-beta-cyclodextrin (MβCD), to disrupt lipid raft-mediated PA internalization. Treatment with MβCD prevented the PA-induced loss of lamellar cristae morphology. Left column, 3000x magnification, scale bar: 1 μm. Right column, 20,000x magnification, scale bar: 200 nm. N=3.

**Figure 10:**
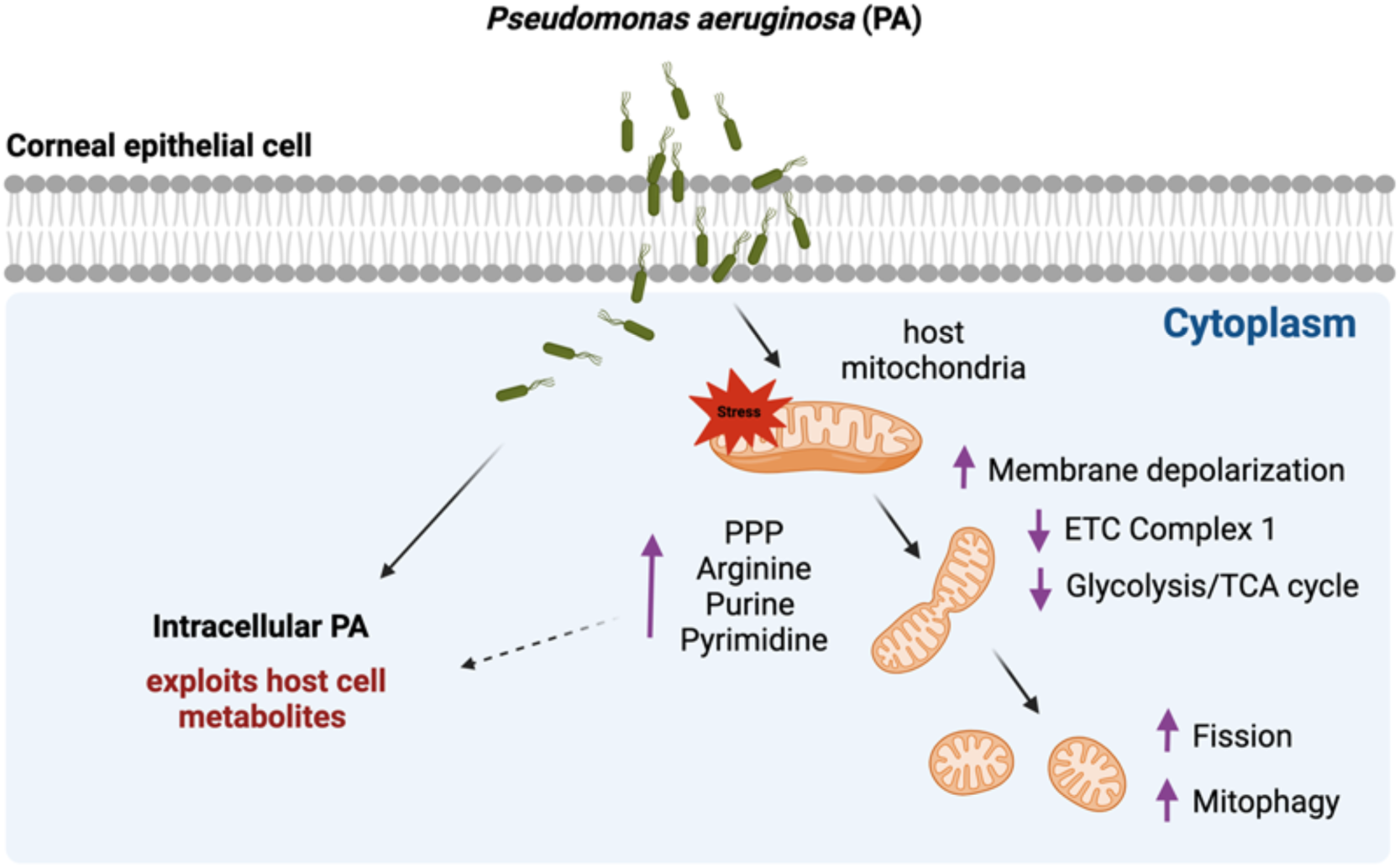
A schematic of PA induced disruption of host cell mitochondria and metabolic rewiring. PA infection of corneal epithelial cells impairs host mitochondria. The increase in the pentose phosphate pathway (PPP), arginine biosynthesis, and purine and pyrimidine metabolism provides intracellular PA with nucleotides, amino acids, and nitrogen (dashed arrow). Mitophagy allows for removal of damaged mitochondria to promote host cell survival.

## Discussion

In the current study, we investigated the effects of PA infection on mitochondrial homeostasis and metabolism in corneal epithelial cells. Our first key finding is that PA infection induces mitochondrial fission and a loss of the lamellar cristae architecture. The increase in phosphorylated DRP1 immediately following infection confirms that mitochondrial fission occurs in a DRP-1 dependent manner. In addition to changes in mitochondrial ultrastructure, PA also induced a loss of mitochondrial membrane potential. Various bacterial virulence factors such as effector proteins and pore forming toxins have been speculated to regulate mitochondrial homeostasis during infection.(34–36) In addition to mitochondrial fragmentation, the T3SS has been shown to induce mitochondrial depolarization and cell apoptosis via GTPase-activating protein (GAP) Domain Activity in HeLa cells.(37) Since Type III effectors are known to act immediately following contact with the host cell, this would explain the rapid induction of fission evidenced in our study.(38)

Consistent with the loss of mitochondrial depolarization, we found an increase in PINK1-mediated mitophagy. During normal homeostasis, PINK1 undergoes rapid turnover by the proteasome.(39) Upon depolarization however, PINK1 accumulates at the outer mitochondrial membrane. This leads to the translocation of Parkin and the subsequent recruitment of the autophagy machinery that is needed for clearance of damaged mitochondria. In addition to PINK1, we also found an increase in FUNDC1-mediated mitophagy. FUNDC1 has been shown to play a role in hypoxia-induced mitophagy and is particularly important in cardiomyocytes.(40) The role of FUNDC1 in mitophagy during infection in less clear. FUNDC1 has been reported to associate at mitochondrial membrane contact sites and may influence mitochondrial tethering to other organelles.(41, 42) FUNDC1 has also been shown to interact with DRP1 and OPA1 to mediate mitochondrial dynamics.(40) Thus, the increase in FUNDC1 may potentiate mitochondrial fission induced by PA. In contrast to PINK1 and FUNDC1, BNIP3L/NIX mediated mitophagy was downregulated. We have previously shown that BNIP3L/NIX mediated mitophagy occurs as part of normal homeostasis in the corneal epithelium, with a gradient pattern of expression.(43) This suggests a role for BNIP3L/NIX in mitophagy during cellular differentiation. While further studies are needed to investigate the role of BNIP3L/NIX in corneal epithelial cell differentiation, the present data indicates that PA induces a switch from physiological mitophagy through BNIP3L/NIX to PINK1 and FUNDC1 pathways. Different types of bacteria have been shown to induce selective mitophagy pathways during infection in host cells.(44) In some instances, the induction of mitophagy has been shown to convey a survival advantage to the host through the inhibition of xenophagy.(45) In other cases, bacteria induce mitophagy to facilitate the removal of damaged mitochondria and limit the production of reactive oxygen species (ROS) that are detrimental to their intracellular survival.(16, 22) Indeed, PA has been shown to induce PINK1-mediated mitophagy in *C. elegans*, where mitophagy conferred protection against pyoverdine, a siderophore released by PA.(23) In macrophages, mitophagy induced via the receptor prohibitin 2 (PHB2) in response to PA infection facilitates bacterial clearance through scavenging ROS.(46) In a related study, macrophages were also shown to exhibit TANK-binding kinase-1 (TBK1)-mediated mitophagy during PA infection.(47) This was due to the presence of host mtDNA and the induction of cGAS, thereby demonstrating a direct link between mitochondrial homeostasis and inflammation.

A major metabolic pathway for most eukaryotic cells is oxidative phosphorylation. Like mitochondrial fission and mitophagy, oxidative phosphorylation is known to be a major target by pathogens.(48) The first step in this pathway is the oxidation of NADH at complex 1. NADH is generated primarily by the TCA cycle and is thought to contribute to approximately 40% of the proton motive energy in the electron transport chain.(49) In macrophages deficient for a protein involved in complex 1 assembly, ECSIT, *Salmonella typhimurium* exhibits enhanced intracellular replication. (50) Both *Escherichia coli* and *Salmonella enterica* have been shown to decrease complex 1 in murine macrophages.(51) In contrast to these studies, *Staphylococcus saprophyticus* infection induces GRIM19 and complex I activity.(52) Here we investigated four of the five protein complexes involved in oxidative phosphorylation. Of these, PA infection downregulated complex 1, similar to other gram-negative bacteria.

An important metric of cell health is spare respiratory capacity (SRC).(53) The SRC relates to the capacity a cell has to respond to stress. In our model, the SRC was decreased by both viable and heat killed PA, although to a much lesser extent in the latter group. The change in SRC in response to heat killed PA suggests that the presence of pattern recognition receptors alone are sufficient to exert negative effects on cell health, despite maintenance of basal oxygen consumption. The 30 minute treatment period with gentamicin was not only able to restore SRC to that of the uninfected control, but was sufficient for restoration of mitochondria. This was reflected in both oxygen consumption and mitochondrial structure. This indicates that extracellular and not intracellular PA induce mitochondrial damage during infection and that mitochondria are able to undergo repair despite the presence of intracellular PA. Even more intriguing however, was the observed increase in SRC following treatment with the lipid raft inhibitor, MβCD. Since MβCD had no toxic effect on PA and has only been shown to reduce intracellular levels of PA by half, this begs the question as to whether MβCD induced membrane changes prevented cristae loss or whether the reduced number of PA that entered the cell facilitated more rapid mitochondrial repair.

Analysis of the metabolite profile supported our findings of impaired mitochondrial function. This was accompanied by a shift towards the pentose phosphate pathway. This shift is likely due in part to the need to counteract PA-induced ROS in host cells.(54) Of all the metabolic pathways, arginine biosynthesis was the most impacted pathway that was increased during PA infection. Within host cells, nitric oxide synthase converts arginine to nitric oxide and citrulline, the former of which exerts antibacterial effects. Indeed, we found a significant increase in citrulline in PA infected cells supporting increased flux through this pathway. Similar to host cells, certain types of bacteria, including PA, are able to metabolize arginine as a source of nitrogen.(55) This raises the possibility of competition between host and pathogen for the arginine pool. Purine and pyrimidine metabolism was also significantly elevated in our model. Like arginine, bacteria are able to utilize host purines and pyrimidines for energy generation. In gut dwelling bacteria, purine degradation has been shown to occur in an anaerobic manner.(56) Pyrimidines also provide PA with added nitrogen to support anaerobic metabolism.(57) Given the observed differences in oxygen consumption between intra- and extracellular PA, together these data suggest that intracellular PA may undergo a metabolic switch and exploit key metabolic pathways in host cells for adequate nutrition.

It is now well established that intracellular PA is able to form membrane blebs in corneal epithelial cells. These blebs, which are devoid of cytoskeletal components, are replicative niches for the bacteria.(58) Prior work has shown that bleb formation is independent of changes in the actin cytoskeleton but is impacted by osmolarity.(5) Interestingly, in our model, we found a significant increase in betaine metabolism. Betaine is a vital osmolyte that functions to protect cells from osmotic stress and promote cell volume.(59) Betaine also functions to buffer mitochondrial ROS and regulate mitochondrial dynamics.(60) Unlike betaine, biotin metabolism was the most downregulated metabolic pathway during infection. Biotin is involved in a myriad of cellular functions and can also be metabolized by bacteria. In *Francisella sp*., biotin is essential for phagosomal escape.(61) This raises the possibility as to whether PA uses host cell biotin to rapidly escape the phagosome, preventing clearance by xenophagy.

In conclusion, we show that PA induces mitochondrial damage as part of the invasion process in corneal epithelial cells. During this time, PA is highly aerobic and host mitochondrial function is impaired. Onside intracellular, PA appears to be undergoing a metabolic shift. More study is needed to elucidate the changes in PA metabolism at a molecular level and to test whether PA mutants defective in anaerobic metabolism are able to internalize. Importantly, this potential metabolic shift in PA corresponds to the restoration of mitochondrial function in host cells. The induction of multiple mitophagy pathways suggests that this mechanism is essential to facilitate mitochondrial recovery and promote host cell survival. Further studies are needed to link these mitochondrial changes with bleb formation and intracellular trafficking of PA, and to determine the long term immunometabolic effects of PA replication and survival in the corneal epithelium.

## References

1. Yamamoto N, Yamamoto N, Petroll MW, Cavanagh HD, Jester JV. Internalization of *Pseudomonas aeruginos*a is mediated by lipid rafts in contact lens-wearing rabbit and cultured human corneal epithelial cells. Invest Ophthalmol Vis Sci. 2005;46(4):1348–55.

2. Zaidi T, Bajmoczi M, Zaidi T, Golan DE, Pier GB. Disruption of CFTR-dependent lipid rafts reduces bacterial levels and corneal disease in a murine model of *Pseudomonas aeruginosa* keratitis. Invest Ophthalmol Vis Sci. 2008;49(3):1000–9.

3. Yamamoto N, Yamamoto N, Jester JV, Petroll WM, Cavanagh HD. Prolonged hypoxia induces lipid raft formation and increases *Pseudomonas* internalization in vivo after contact lens wear and lid closure. Eye Contact Lens. 2006;32(3):114–20.

4. Yamamoto N, Yamamoto N, Petroll MW, Jester JV, Cavanagh HD. Regulation of *Pseudomonas aeruginosa* internalization after contact lens wear in vivo and in serum-free culture by ocular surface cells. Invest Ophthalmol Vis Sci. 2006;47(8):3430–40.

5. Jolly AL, Takawira D, Oke OO, Whiteside SA, Chang SW, Wen ER, et al. *Pseudomonas aeruginosa*-induced bleb-niche formation in epithelial cells is independent of actinomyosin contraction and enhanced by loss of cystic fibrosis transmembrane-conductance regulator osmoregulatory function. mBio. 2015;6(2):e02533.

6. Angus AA, Evans DJ, Barbieri JT, Fleiszig SM. The ADP-ribosylation domain of *Pseudomonas aeruginosa* ExoS is required for membrane bleb niche formation and bacterial survival within epithelial cells. Infect Immun. 2010;78(11):4500–10.

7. Angus AA, Lee AA, Augustin DK, Lee EJ, Evans DJ, Fleiszig SM. *Pseudomonas aeruginosa* induces membrane blebs in epithelial cells, which are utilized as a niche for intracellular replication and motility. Infect Immun. 2008;76(5):1992–2001.

8. Hritonenko V, Mun JJ, Tam C, Simon NC, Barbieri JT, Evans DJ, et al. Adenylate cyclase activity of *Pseudomonas aeruginosa* ExoY can mediate bleb-niche formation in epithelial cells and contributes to virulence. Microb Pathog. 2011;51(5):305–12.

9. Andrieux P, Chevillard C, Cunha-Neto E, Nunes JPS. Mitochondria as a Cellular Hub in Infection and Inflammation. Int J Mol Sci. 2021;22(21).

10. Nunnari J, Suomalainen A. Mitochondria: in sickness and in health. Cell. 2012;148(6):1145–59.

11. Vringer E, Tait SWG. Mitochondria and cell death-associated inflammation. Cell Death Differ. 2023;30(2):304–12.

12. Shen K, Pender CL, Bar-Ziv R, Zhang H, Wickham K, Willey E, et al. Mitochondria as Cellular and Organismal Signaling Hubs. Annu Rev Cell Dev Biol. 2022;38:179–218.

13. Bravo-Sagua R, Parra V, López-Crisosto C, Díaz P, Quest AF, Lavandero S. Calcium Transport and Signaling in Mitochondria. Compr Physiol. 2017;7(2):623–34.

14. Ashrafi G, Schwarz TL. The pathways of mitophagy for quality control and clearance of mitochondria. Cell Death Differ. 2013;20(1):31–42.

15. Tiku V, Tan MW, Dikic I. Mitochondrial Functions in Infection and Immunity. Trends Cell Biol. 2020;30(4):263–75.

16. Zhang Y, Yao Y, Qiu X, Wang G, Hu Z, Chen S, et al. *Listeria* hijacks host mitophagy through a novel mitophagy receptor to evade killing. Nat Immunol. 2019;20(4):433–46.

17. Lum M, Morona R. Dynamin-related protein Drp1 and mitochondria are important for Shigella flexneri infection. Int J Med Microbiol. 2014;304(5-6):530–41.

18. Escoll P, Song OR, Viana F, Steiner B, Lagache T, Olivo-Marin JC, et al. *Legionella pneumophila* Modulates Mitochondrial Dynamics to Trigger Metabolic Repurposing of Infected Macrophages. Cell Host Microbe. 2017;22(3):302–16.e7.

19. Emelyanov VV. Mitochondrial porin VDAC 1 seems to be functional in rickettsial cells. Ann N Y Acad Sci. 2009;1166:38–48.

20. Chowdhury SR, Reimer A, Sharan M, Kozjak-Pavlovic V, Eulalio A, Prusty BK, et al. *Chlamydia* preserves the mitochondrial network necessary for replication via microRNA-dependent inhibition of fission. J Cell Biol. 2017;216(4):1071–89.

21. Mahla RS, Kumar A, Tutill HJ, Krishnaji ST, Sathyamoorthy B, Noursadeghi M, et al. NIX-mediated mitophagy regulate metabolic reprogramming in phagocytic cells during mycobacterial infection. Tuberculosis (Edinb). 2021;126:102046.

22. Jiao Y, Cao S, Zhang Y, Tan Y, Zhou Y, Wang T, et al. *Yersinia pestis*-Induced Mitophagy That Balances Mitochondrial Homeostasis and mROS-Mediated Bactericidal Activity. Microbiol Spectr. 2022;10(3):e0071822.

23. Kirienko NV, Ausubel FM, Ruvkun G. Mitophagy confers resistance to siderophore-mediated killing by *Pseudomonas aeruginosa*. Proc Natl Acad Sci U S A. 2015;112(6):1821–6.

24. Jabir MS, Hopkins L, Ritchie ND, Ullah I, Bayes HK, Li D, et al. Mitochondrial damage contributes to *Pseudomonas aeruginosa* activation of the inflammasome and is downregulated by autophagy. Autophagy. 2015;11(1):166–82.

25. Maurice NM, Bedi B, Yuan Z, Goldberg JB, Koval M, Hart CM, et al. *Pseudomonas aeruginosa* Induced Host Epithelial Cell Mitochondrial Dysfunction. Sci Rep. 2019;9(1):11929.

26. Josephson H, Ntzouni M, Skoglund C, Linder S, Turkina MV, Vikström E. *Pseudomonas aeruginosa* N-3-Oxo-Dodecanoyl-Homoserine Lactone Impacts Mitochondrial Networks Morphology, Energetics, and Proteome in Host Cells. Front Microbiol. 2020;11:1069.

27. Tounta V, Liu Y, Cheyne A, Larrouy-Maumus G. Metabolomics in infectious diseases and drug discovery. Mol Omics. 2021;17(3):376–93.

28. Russell DG, Huang L, VanderVen BC. Immunometabolism at the interface between macrophages and pathogens. Nat Rev Immunol. 2019;19(5):291–304.

29. Riquelme SA, Prince A. *Pseudomonas aeruginosa* Consumption of Airway Metabolites Promotes Lung Infection. Pathogens. 2021;10(8).

30. Riquelme SA, Liimatta K, Wong Fok Lung T, Fields B, Ahn D, Chen D, et al. *Pseudomonas aeruginosa* Utilizes Host-Derived Itaconate to Redirect Its Metabolism to Promote Biofilm Formation. Cell Metab. 2020;31(6):1091–106.e6.

31. Robertson DM, Li L, Fisher S, Pearce VP, Shay JW, Wright WE, et al. Characterization of Growth and Differentiation in a Telomerase-Immortalized Human Corneal Epithelial Cell Line. Invest Ophthalmol Vis Sci. 2005;46(2):470–8.

32. Schneider CA, Rasband WS, Eliceiri KW. NIH Image to ImageJ: 25 years of image analysis. Nat Methods. 2012;9(7):671–5.

33. Yuan M, Breitkopf SB, Yang X, Asara JM. A positive/negative ion–switching, targeted mass spectrometry–based metabolomics platform for bodily fluids, cells, and fresh and fixed tissue. Nature Protocols. 2012;7(5):872–81.

34. Stavru F, Bouillaud F, Sartori A, Ricquier D, Cossart P. *Listeria monocytogenes* transiently alters mitochondrial dynamics during infection. Proc Natl Acad Sci U S A. 2011;108(9):3612–7.

35. Foo JH, Culvenor JG, Ferrero RL, Kwok T, Lithgow T, Gabriel K. Both the p33 and p55 subunits of the *Helicobacter pylori* VacA toxin are targeted to mammalian mitochondria. J Mol Biol. 2010;401(5):792–8.

36. Fine-Coulson K, Giguère S, Quinn FD, Reaves BJ. Infection of A549 human type II epithelial cells with *Mycobacterium tuberculosis* induces changes in mitochondrial morphology, distribution and mass that are dependent on the early secreted antigen, ESAT-6. Microbes and Infection. 2015;17(10):689–97.

37. Wood SJ, Goldufsky JW, Bello D, Masood S, Shafikhani SH. *Pseudomonas aeruginosa* ExoT Induces Mitochondrial Apoptosis in Target Host Cells in a Manner That Depends on Its GTPase-activating Protein (GAP) Domain Activity*. Journal of Biological Chemistry. 2015;290(48):29063–73.

38. Lee VT, Smith RS, Tümmler B, Lory S. Activities of *Pseudomonas aeruginosa* effectors secreted by the Type III secretion system in vitro and during infection. Infect Immun. 2005;73(3):1695–705.

39. Jin SM, Youle RJ. PINK1- and Parkin-mediated mitophagy at a glance. J Cell Sci. 2012;125(Pt 4):795–9.

40. Chen M, Chen Z, Wang Y, Tan Z, Zhu C, Li Y, et al. Mitophagy receptor FUNDC1 regulates mitochondrial dynamics and mitophagy. Autophagy. 2016;12(4):689–702.

41. Wu W, Lin C, Wu K, Jiang L, Wang X, Li W, et al. FUNDC1 regulates mitochondrial dynamics at the ER-mitochondrial contact site under hypoxic conditions. Embo j. 2016;35(13):1368–84.

42. Chai P, Cheng Y, Hou C, Yin L, Zhang D, Hu Y, et al. USP19 promotes hypoxia-induced mitochondrial division via FUNDC1 at ER-mitochondria contact sites. J Cell Biol. 2021;220(7).

43. Stuard Sambhariya W, Trautmann IJ, Robertson DM. Insulin-like growth factor binding protein-3 mediates hyperosmolar stress-induced mitophagy through the mechanistic target of rapamycin. J Biol Chem. 2023;299(11):105239.

44. Verbeke J, De Bolle X, Arnould T. To eat or not to eat mitochondria? How do host cells cope with mitophagy upon bacterial infection? PLoS Pathog. 2023;19(7):e1011471.

45. Song Y, Ge X, Chen Y, Hussain T, Liang Z, Dong Y, et al. *Mycobacterium bovis* induces mitophagy to suppress host xenophagy for its intracellular survival. Autophagy. 2022;18(6):1401–15.

46. Huang T, Pu Q, Zhou C, Lin P, Gao P, Zhang X, et al. MicroRNA-302/367 Cluster Impacts Host Antimicrobial Defense via Regulation of Mitophagic Response Against *Pseudomonas aeruginosa* Infection. Front Immunol. 2020;11:569173.

47. Wang B, Zhou C, Wu Q, Lin P, Pu Q, Qin S, et al. cGAS modulates cytokine secretion and bacterial burdens by altering the release of mitochondrial DNA in *Pseudomonas* pulmonary infection. Immunology. 2022;166(3):408–23.

48. Escoll P, Buchrieser C. Metabolic reprogramming of host cells upon bacterial infection: Why shift to a Warburg-like metabolism? Febs j. 2018;285(12):2146–60.

49. Escoll P, Platon L, Buchrieser C. Roles of Mitochondrial Respiratory Complexes during Infection. Immunometabolism. 2019;1(2):e190011.

50. West AP, Brodsky IE, Rahner C, Woo DK, Erdjument-Bromage H, Tempst P, et al. TLR signalling augments macrophage bactericidal activity through mitochondrial ROS. Nature. 2011;472(7344):476–80.

51. Garaude J, Acín-Pérez R, Martínez-Cano S, Enamorado M, Ugolini M, Nistal-Villán E, et al. Mitochondrial respiratory-chain adaptations in macrophages contribute to antibacterial host defense. Nat Immunol. 2016;17(9):1037–45.

52. Chen Y, Lu H, Liu Q, Huang G, Lim CP, Zhang L, et al. Function of GRIM-19, a mitochondrial respiratory chain complex I protein, in innate immunity. J Biol Chem. 2012;287(32):27227–35.

53. Marchetti P, Fovez Q, Germain N, Khamari R, Kluza J. Mitochondrial spare respiratory capacity: mechanisms, regulation, and significance in non-transformed and cancer cells. FASEB J. 2020;34(10):13106–24.

54. Liu CL, Hsu YC, Lee JJ, Chen MJ, Lin CH, Huang SY, et al. Targeting the pentose phosphate pathway increases reactive oxygen species and induces apoptosis in thyroid cancer cells. Mol Cell Endocrinol. 2020;499:110595.

55. Luckett JC, Darch O, Watters C, Abuoun M, Wright V, Paredes-Osses E, et al. A novel virulence strategy for *Pseudomonas aeruginosa* mediated by an autotransporter with arginine-specific aminopeptidase activity. PLoS Pathog. 2012;8(8):e1002854.

56. Kasahara K, Kerby RL, Zhang Q, Pradhan M, Mehrabian M, Lusis AJ, et al. Gut bacterial metabolism contributes to host global purine homeostasis. Cell Host Microbe. 2023;31(6):1038–53.e10.

57. Garavito MF, Narváez-Ortiz HY, Zimmermann BH. Pyrimidine Metabolism: Dynamic and Versatile Pathways in Pathogens and Cellular Development. J Genet Genomics. 2015;42(5):195–205.

58. Heimer SR, Evans DJ, Stern ME, Barbieri JT, Yahr T, Fleiszig SM. *Pseudomonas aeruginosa* utilizes the type III secreted toxin ExoS to avoid acidified compartments within epithelial cells. PLoS One. 2013;8(9):e73111.

59. Garrett Q, Khandekar N, Shih S, Flanagan JL, Simmons P, Vehige J, et al. Betaine stabilizes cell volume and protects against apoptosis in human corneal epithelial cells under hyperosmotic stress. Exp Eye Res. 2013;108:33–41.

60. Jung Kim M. Betaine enhances the cellular survival via mitochondrial fusion and fission factors, MFN2 and DRP1. Anim Cells Syst (Seoul). 2018;22(5):289–98.

61. Napier BA, Meyer L, Bina JE, Miller MA, Sjöstedt A, Weiss DS. Link between intraphagosomal biotin and rapid phagosomal escape in Francisella. Proc Natl Acad Sci U S A. 2012;109(44):18084–9.

