## Supporting data for "*Pseudomonas aeruginosa* impairs mitochondrial function and metabolism during infection of corneal epithelial cells"

**Supporting Figures**

**Supp. Figure 1: PA infection induces mitochondrial depolarization in primary cultured corneal epithelial cells.** Primary corneal epithelial cells (HCEC) were infected with PAO1 for 2 hours and then labeled with JC-1. Similar to hTCEpi cells, PA infection resulted in robust mitochondrial depolarization. Scale bar: 21.2 μm.

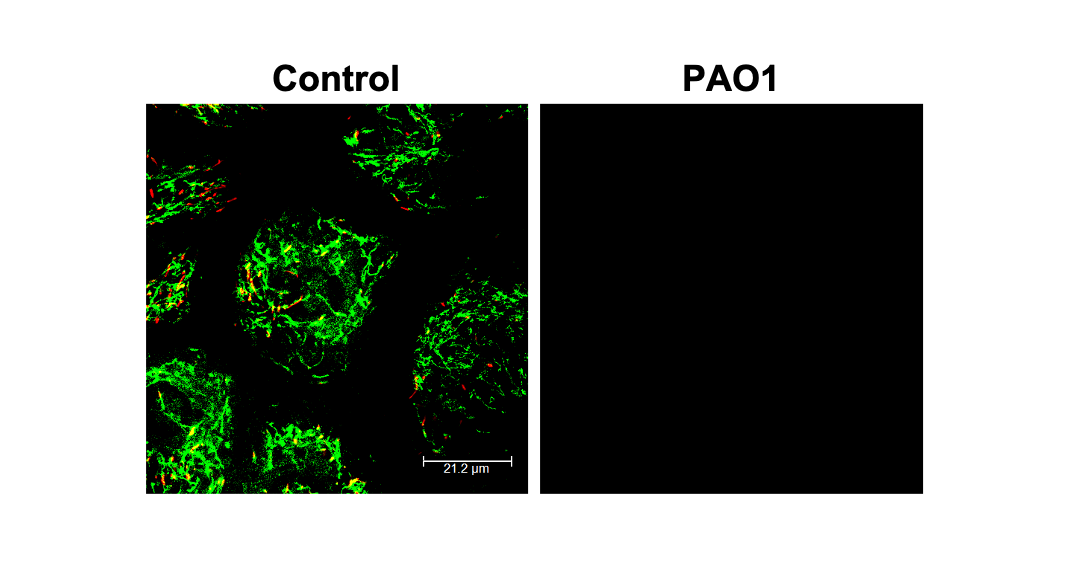

**Supp. Figure 2:** **Pathway enrichment of the altered metabolites during PA infection.** (A) Enrichment overview of the top 25 pathways detected from 34 significant metabolites between groups. (B) Enrichment overview of the top 25 pathways detected from 26 significantly upregulated metabolites in the PA infected group. (C) Enrichment overview of the top 25 pathways related to the 8 significantly downregulated metabolites in the PA infected group. Untargeted metabolomics N=5, MetaboAnalyst 5.0.

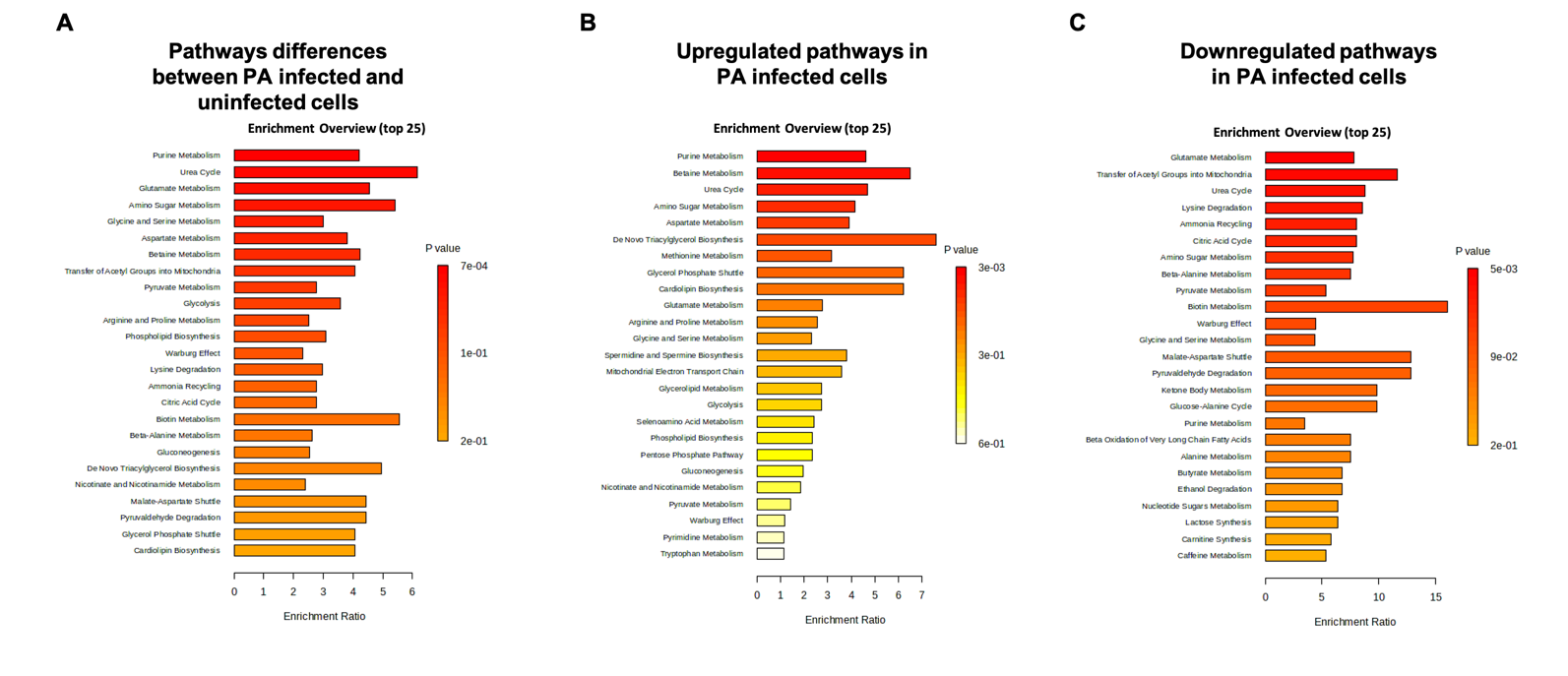

**Supp. Figure 3: PA intracellular survival during gentamicin treatment.** hTCEpi cells were infected with PAO1 for 2 hours, followed by treatment with 200 µg/mL gentamicin for 1 hour. PBS only represents PA infected cells in PBS without gentamicin. Data are normalized to the PBS only condition, N=3, **p<0.01, students t-test.

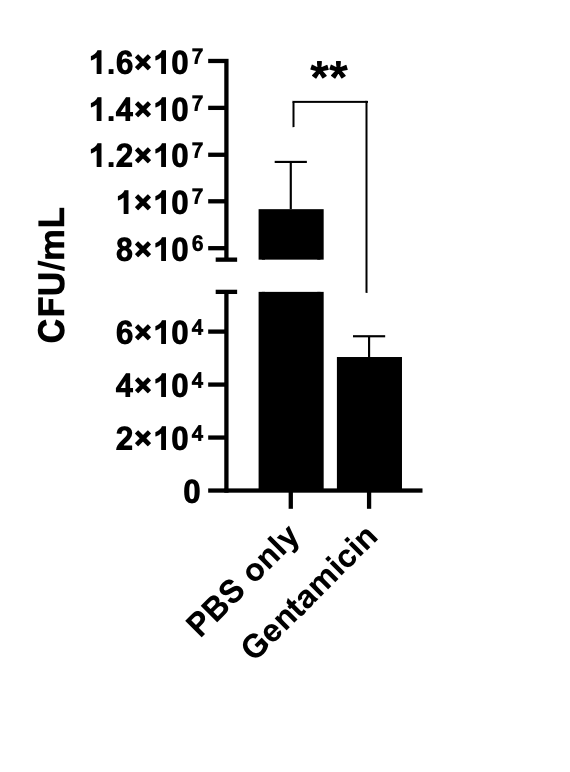
